# The BumbleBox: An open-source platform for quantifying behavior in bumblebee colonies

**DOI:** 10.1101/2024.11.07.622358

**Authors:** August Easton-Calabria, James D. Crall

## Abstract

1. Bumblebees (Apini: *Bombus*) are important pollinators globally and an emerging model system for studying the ecology and evolution of social behavior and effects of environmental stressors on bees.
2. Behavioral studies of bumblebees have conventionally relied on labor and time-intensive manual observations. While recent years have seen rapid advances in automated behavioral tracking in social insects, these tracking technologies are often expensive and require extensive programming experience, limiting accessibility and widespread adoption.
3. Here we introduce the BumbleBox, an open-source system for automated tracking and behavioral quantification of individual bumblebees that can be built using low-cost consumer components and DIY fabrication (i.e., 3D-printing and laser-cutting). We provide an integrated pipeline for data collection and analysis, including nest arena design, software for automated collection of video data, and the quantification of individual behavior.
4. The BumbleBox system is designed to be (a) ***accessible***, requiring no prior experience with programming or hardware design to operate; (b) ***scalable***, allowing long-term, automated tracking across many units in parallel at low-cost; and (c) ***modular***, allowing for flexible adoption to unique applications in bumblebees and other systems. We validate the use of this system in a widespread bumblebee species (*Bombus impatiens*) that is both commercially and ecologically important. Finally, we highlight widespread potential applications in quantifying behavior and pollinator health in bumblebees and other social insects, including screening impacts of pesticides and other environmental stressors on social behavior.

## 1. Introduction

Social bees are critically important for the functioning of ecosystems and the delivery of pollination services across the globe (Klein et al., 2006). Bumblebees (*Apini: Bombus*) are a diverse group of generalist pollinators that are among the most abundant pollinators globally (∼260 species (Cameron & Sadd, 2020)). Bumblebees are economically important for crop pollination (Fijen et al., 2018) and an emerging model system for studying social behavior and the effects of environmental stressors on bees (Woodard et al., 2015) (Easton-Calabria et al., 2023) (Amsalem et al., 2014) (Baer, 2003) (Gill et al., 2012).

Automated tracking and behavioral quantification within social insect colonies has advanced rapidly in recent years (Mersch et al., 2013) (Crall, Switzer, et al., 2018) (Boenisch et al., 2018) (Sclocco et al., 2021), providing fundamental insight into social interaction networks (Richardson et al., 2017), disease dynamics (Stroeymeyt et al., 2018), and effects of anthropogenic stress (Easton-Calabria et al., 2023). However, methods for automated tracking have been expensive, computationally intensive, and require substantial experience in hardware design and computer programming. The emergence of open-source, inexpensive, and low-power electronics, however, are making scalable quantification of social behavior feasible and accessible; this creates new opportunities for standardized, reproducible methods of automated behavioral quantification to address key knowledge gaps in bumblebee behavior, ecology, and health.

Here, we introduce the BumbleBox, an automated, open-source hardware and software imaging system for automated tracking and behavioral quantification of uniquely identified bumblebees within the nest. The BumbleBox system is designed to be: (a) accessible, including open-source design and low-cost, (b) scalable, with capacity for parallel, automated data analysis across multiple units and extended time periods, and (c) modular, with adaptability for different experimental contexts and applications. We first describe the basic structure, function, and operation of the system, then describe the integrated pipeline from arena design and experimental collection of video data to the quantification of individual behavior. Finally, we demonstrate the application of this system for automated tracking of individual behavior within bumblebee colonies, and explore its use in different experimental contexts and applications (e.g., across multiple species in the lab and field).

## 2. Methods

Here, we provide a basic overview of the BumbleBox operation and capacity. The BumbleBox is designed to provide automation of experimental tasks such as video recording, tracking, and behavioral quantification within bumblebee colonies on the Raspberry Pi (Saunders et al., 2022). A primary goal of this system is accessibility for researchers and practitioners across fields and a detailed guide for construction, installation, and use is provided at https://github.com/Crall-Lab/BumbleBox.

### 2.1 Design and hardware

The BumbleBox consists of a nest arena and imaging module for automated recording (Fig 1a). The nest arena (154×154×275 mm) houses the bumblebee (*Bombus* spp) colony, allows for viewing and feeding of the colony, and accommodates imaging under controlled infrared lighting conditions. It is fabricated from 3D printed and laser cut components because of (a) the increasing availability of these fabrication techniques (e.g., open-access maker spaces or online fabrication services), (b) cost-effectiveness (total material costs: ∼$350), and (c) modularity and customizability.

**Figure 1.**
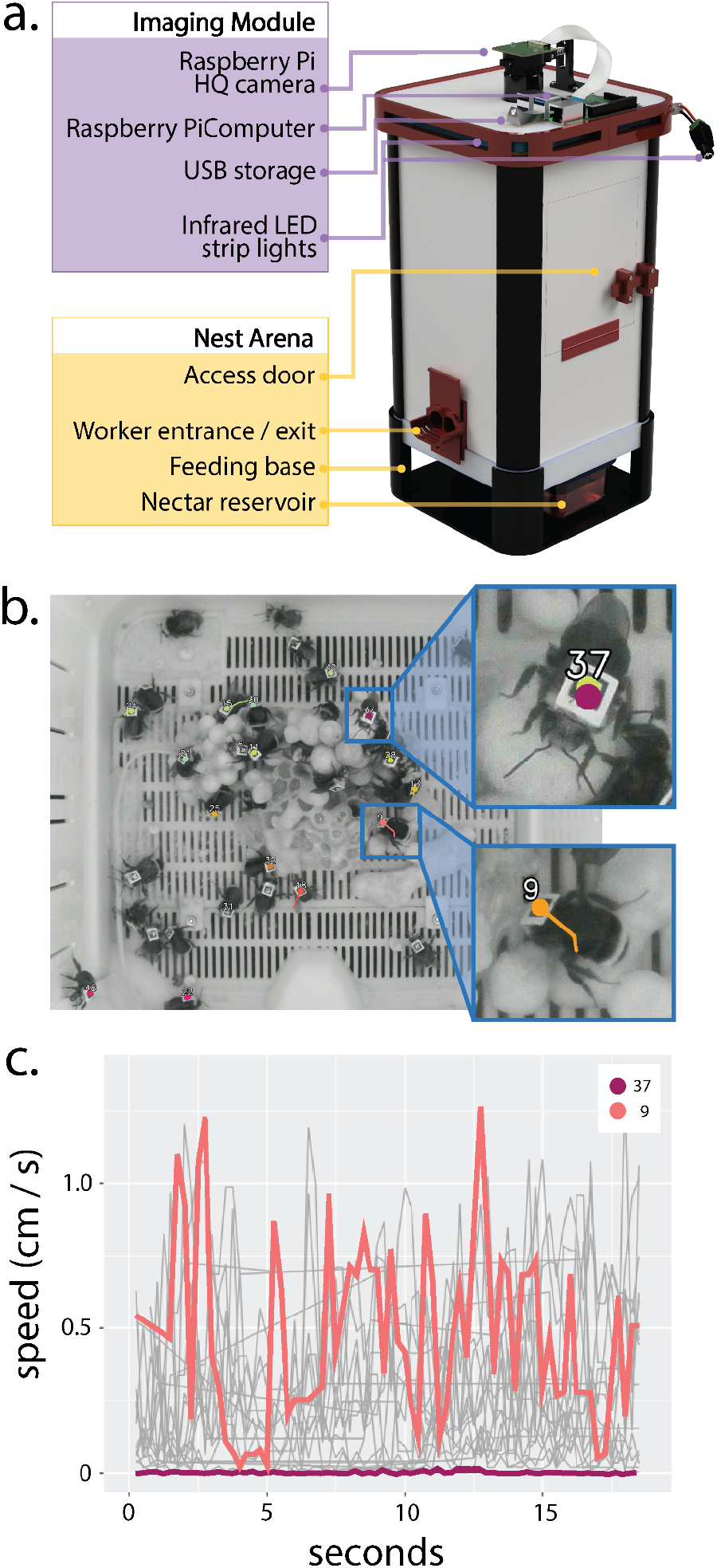
Visualization of the BumbleBox tracking systems and example data output. a) A 3D-render of the custom BumbleBox design. b) Example tracking within the BumbleBox, with insets showing two example Individuals with locations tracked using ArUco tags. c) The speeds of two individuals (same individuals as in (b)) plotted over the course of a single 20 second recording period.

Imaging within BumbleBoxes occurs *via* the Raspberry Pi high-quality camera (RPi HQ, 12 megapixels, 4056×3040 pixels), which is mounted on top of the device (Fig 1a) and provides recording of the colony viewed from above. Within the nest, illumination is provided by infrared LEDs (840 nm), as bees cannot perceive near-infrared light (Martínez-Harms et al., 2010). To provide imaging in the near-IR spectrum, the RPi HQ camera is modified to remove the IR-blocking filter.

### 2.2 Individual bee tracking

The BumbleBox uses ArUco (an OpenCV module, implemented in Python) markers to capture the position of individual bees within the nest - ArUco marker detection software is open-source, community-maintained, and the markers can be printed on paper and detected at small sizes within images (Garrido-Jurado et al., 2014). The ArUco software tracks tags from predefined tag dictionaries, and offers dictionaries of different marker sizes and tag number (ranging from 50-1,000 tags). The BumbleBox Github repository includes high-quality image files of each standard tag dictionary for printing, and allows users to easily change between dictionaries being used for tracking.

### 2.3 Experimental workflow

The BumbleBox software supports: (1) setup, (2) data acquisition, and (3) behavioral analysis. The BumbleBox uses custom Python scripts to automate data capture and analyze behavioral data but previous knowledge of Python is not required. For example, users start collecting data by customizing a single script (setup.py) (Table 1). The default recording values have been tested using the standard nest arena designs for the BumbleBox described here (imaged with the Raspberry Pi High Quality Camera, and infrared 840nm LEDs). Users then start the automation of data capture by simply running a single script (start_automated_recording.py).

**Table 1.** Adjustable recording parameters for the BumbleBox in the setup.py script.

| Parameter name | Description |
| --- | --- |
| Recording length (seconds) | The length of individual recording periods. Default: 20 seconds |
| Tag tracking (True/False) | Track Aruco tags from frames immediately after capture and store tag data in a text file, without saving a video |
| Data capture frequency (once every X minutes) | Choose how often data is captured. Possible values: 1-10, 15, 20, 30, 60 |
| Video recording frequency (once every X minutes) | Choose how often videos are saved. Overridden by data capture frequency if tag tracking is turned off (every recording would be saved as a video) |
| Video width, height (pixels) | Choose the video size in pixels. Default: 4056 x 3040 (the Rpi HQ camera maximum resolution) |
| Video file type (MP4 or MJPEG) | Higher framerate can be achieved with the MP4 file type<br>Default: MP4 |
| Shutter speed (microseconds) | Default: 2,500 |
| Frame rate (1 / X frames per second) | Up to 10 frames per second is possible for short videos. Default: 6 |
| Video quality (for MJPEG videos only) | Video quality vs. video size (MJPEG compression). Possible values: 0-100.<br>Default: 95 |
| Infrared recording (True/False) | If true, frames will be processed using an algorithm that accounts for IR light |
| Noise reduction mode | Choose amount of digital noise reduction. Possible values: Auto, None, Fast, High Quality. Default: Auto |
| Digital zoom | Record from a smaller portion of the camera's sensor. Default: None |
| Aruco dictionary | Select the Aruco dictionary that is being used. Default: 4x4_50 |
| Aruco detection parameters | Rather than setting Aruco detection parameters individually, choose to use a preselected set of detection parameters tailored to either the custom or the Koppert-adapted BumbleBox. Default: custom |

#### 2.3.1. Setup

Before initiating data collection, users can ensure optimal imaging conditions (i.e., confirm the nest arena is well lit and tags are visible and in focus) using a script (rpi4_preview.py) that provides a preview of the nest floor. Users can then customize data collection parameters in the

setup script (setup.py), including recording time, frame rate, and whether automatic behavioral quantification should take place after each video (see Table 1 for a description of recording parameters and Table 2 for a description of available behavioral metrics). Then the test tracking script (test_tracking.py) can be run to confirm tags are being read accurately. Once the preview footage and tracking looks correct, the user can begin collecting data by running the automated recording script (start_automated_recording.py, also described above). Data collection begins automatically and will continue whenever the Raspberry Pi is powered.

**Table 2.** Behavioral metrics currently calculated by the BumbleBox software.

| Metric | Description |
| --- | --- |
| Speed | A bee's speed (when moving) calculated across tracked frames, as pixels per second |
| Activity | The proportion of time a bee spends moving vs. at rest per recording period |
| Distance from social center | Each tracked bee's distance from the average x,y position of all tracked bees in a given frame |
| Distance to other bees | Each tracked bee's distance to every other tracked bee in a given frame |
| Contacts | Records contacts between tracked bees in a frame based on a threshold contact distance |
| Distance to nest components | The distance of each tracked bee in a frame to each labelled nest component |

#### 2.3.2 Data Acquisition and Storage

The main function of data acquisition in the BumbleBox is to capture video of the nest arena, run ArUco detection on video frames to detect the location of tagged bees, and to save video and tracking data. Additionally, composite images of the nest without bees (using median background estimation) are stored for later labeling of different nest components, which are useful for a variety of downstream data analyses (e.g., brood care rates).

The user decides how often data acquisition occurs via the setup.py script. There are two types of data outputs: 1) CSV files containing tag coordinate data and behavioral quantification and 2) video files. Each row of the CSV file after tracking contains the detected tag ID, the frame of the recording it was detected within, and the x,y pixel coordinates of the tag center. This is the raw tracking file, and an updated file is generated from it that interpolates missing data and stores the behavioral metrics of interest (see Behavioral Quantification below).

Available data storage can become limited as the generated video files can be large (e.g. ∼40MB for a full-resolution 20 second video at 4fps). Tracking occurs immediately after recording, which allows users to discard videos to reduce the number of videos saved. While it is possible to collect data using the BumbleBox without saving videos at all, saving periodic video files allow a user to check for errors, assess tracking quality, and are useful for manual ground-truth observations and additional downstream analyses (such as pose-tracking (Smith et al., 2022) (Wolf et al., 2023)). Since tags are tracked with and without storing the video data, the user can set different frequencies for tag tracking with and without video recording depending on experimental needs.

#### 2.3.3 Behavioral Quantification

The locations of tagged bees within the nest is used to quantify behavior of uniquely tracked individuals (Table 2). For example, position within the nest, speed, movement, proximity to other bees and nest components, and interactions between individuals can be used to quantify task performance and division of labor within bumblebee colonies (Fig 2, (Crall, Gravish, et al., 2018). Behavioral quantification can take place automatically after instances of data capture or can be calculated on previously collected data, depending on parameters defined in the setup script. While core functions for behavioral quantification are included here, new functions can be added relatively easily for experienced users, and more behavior functions will be added over time. Users can turn on automatic behavior quantification and choose which metrics to be calculated in the setup script (setup.py).

**Figure 2.**
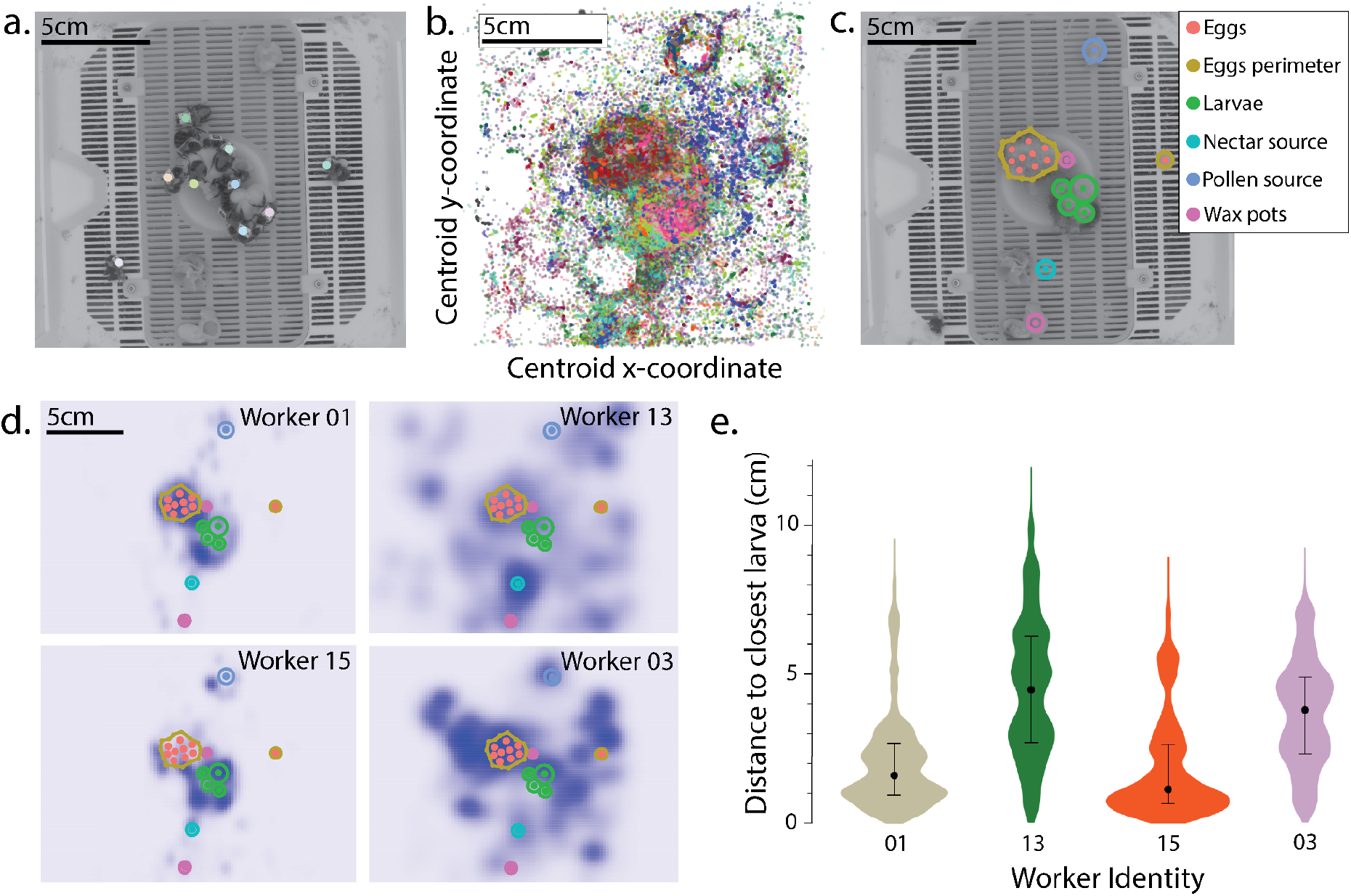
Example quantification of behavior in the BumbleBox. a) Image of a tracked *Bombus impatiens* micracolony from the BumbleBox camera. Solid makers show locations of individually tracked bees. b) Plot showing location data for the microcolony over the course of a single day; each color represents a separate individual. c) Different nest components plotted over a composite nest image generated using the graphical user interface available with the BumbleBox software. d) 2-D gaussian kernel density estimation plots demonstrating the spatial distribution probability of four different individuals within the nest over the course of a single day, with the nest component data overlaid. f) Violin plots depicting probability distributions of each bee’s distance to the closest larva over the course of a single day. Black points Indicate median values and vertical lines represent the interquartile range.

Location and movement of bees relative to nest structure (e.g. brood and wax pots) can be informative for understanding within-nest behavior of bumblebees (Jandt & Dornhaus, 2009) (Crall, Gravish, et al., 2018). The BumbleBox software includes a GUI-based script to label nest structures (e.g., larvae, pupae, wax pots, etc.) and other important features like food sources. Background images for labeling are first generated by median background estimation as in (Crall, Gravish, et al., 2018) (Fig 2c). After labeling, the resulting nest maps can be used to quantify additional behavioral metrics such as minimum distance to wax pots or brood (Fig 2e).

## 3. Results

### 3.1 Validation

We created a microcolony by tagging 16 bumblebee (*Bombus impatiens*) workers with ArUco tags (2.6mm side lengths, printed on TerraSlate paper using a laserjet printer) and imaged them using the BumbleBox system over a period of 24 hours (Fig 2). We briefly anesthetized bees with carbon dioxide and used cyanoacrylate gel superglue to adhere the tags to the thoraxes of the bumblebees. Bees were allowed to recover for 48 hrs before being tracked over a 24-hour period. 240 videos (10 seconds long, ∼5 frames per second, 12,464 total frames) were recorded and 107,242 bee positions were captured. The bees were identified using tags 0-15 (out of a 50 tag

dictionary, 0-49), but there were 7 instances of tracked tags that were among the 34 tags not used (ID 17: 3 instances, ID 26: 1 instance, ID 34: 1 instance, ID 37: 2 instances), and thus considered false positives. Assuming an equivalent false positive rate across tags, we can extrapolate to estimate the number of false positives occurring within tags 0-15 across this recording period: ∼3.29 false positives (0.003%), or an estimated accuracy of >99.99%. An average of 8.6 tags were tracked per frame (median: 9, minimum: 3, lower-quartile: 7, upper-quartile: 10, maximum: 14). See Fig S3 and Supplementary Text for further discussion of tracking performance.

### 3.2 Example applications and modularity

The modularity and scalability of the BumbleBox system allows for diverse applications in the study of bumblebee behavior and health. For example, the BumbleBox system can be adapted for use in field or semi-field conditions: a recent greenhouse experiment modified the BumbleBox to include thermocouples (to quantify brood temperature), as well as the addition of an automated foraging monitoring system (Fig 3a-c, unpublished data). The BumbleBox system is also scalable: a version of the BumbleBox system was used to study the combined effects of thermal stress and neonicotinoid exposure in *B. impatiens* microcolonies (Easton-Calabria et al., 2023), using twelve units in parallel in the lab (Fig 3d). Finally, the BumbleBox has been used across multiple species of *Bombus*, including *B. impatiens, B. griseocollis*, and *B. bimaculatus* from Wisconsin, USA, *B. vosnesenskii* from Oregon, USA, and *B. ephippiatus* (Fig 3e) from Chiapas, Mexico.

**Figure 3.**
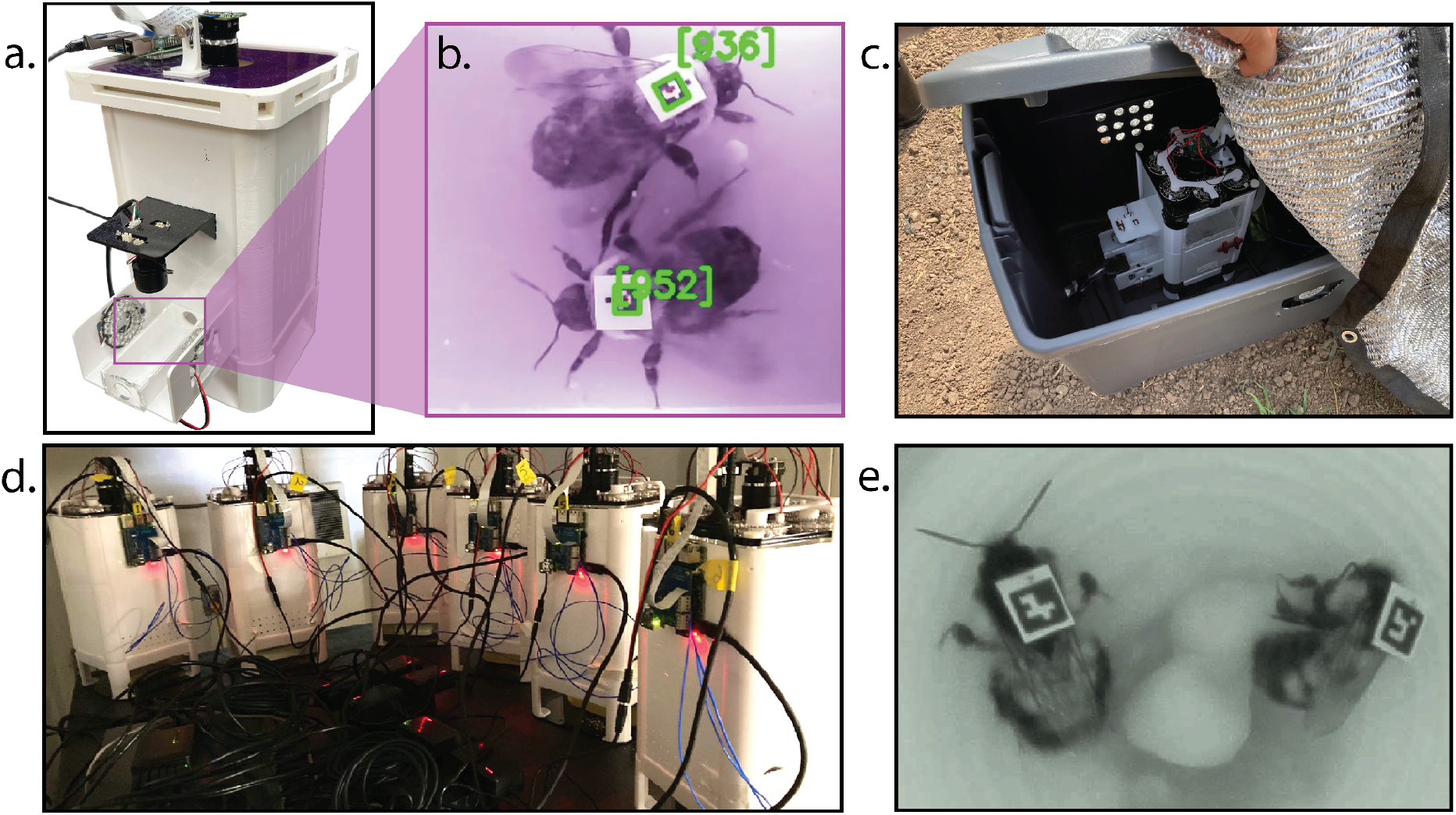
Examples of modular experimental applications of the BumbleBox system. (A) BumbleBox with one wall panel modified as a forage tunnel and outfitted with a second camera (USB) to track foraging activity (B). (C) Modified BumbleBox deployed in the field to monitor foraging activity and in-nest behavior of free-foraging *Bombus impatiens* colonies. (D) Multiple (n = 6) BumbleBoxes outfitted with thermocouples to quantify behavioral responses to thermal stress. (E) A close-up image of two *Bombus ephippiatus* workers from behavioral experiments conducted by collaborators in Chiapas, Mexico.

The design also (optionally) allows users to move the nest from a separate, smaller rearing chamber (145×76×50 mm) into the floor of the nest arena (Supplement Fig 1, design files available at https://github.com/Crall-Lab/BumbleBox). This integration is useful for experiments using microcolonies that need to be established (Klinger et al., 2019), or long-term tracking of colonies reared from wild-caught *Bombus* queens throughout the entire colony cycle (Rowe et al., 2023) (Christman et al., 2023).

An additional BumbleBox nest arena was designed for use with commercially-available (Koppert) bumblebee colony boxes. This design replaces the top of the Koppert box and raises it in order to image the colony from above (Supplementary Fig 2). This was designed with the intention of tagging and imaging mature Koppert commercial colonies, with applications for both lab and field studies.

## 4. Discussion

The BumbleBox is a low-cost, open-source, and modular system for quantifying behavior of uniquely tracked individuals within a bumblebee colony. The system is designed specifically with accessibility to researchers in mind. In particular, it is designed to be usable without modification by users with limited prior knowledge in computer programming or hardware design, while providing flexibility and modularity for experienced users, and has broad research applications in bumblebees and potentially other social insect species (Guo et al., 2024).

The standardized, automated behavioral tracking provided by the BumbleBox has diverse potential applications in bumblebee behavior, ecology, and health. First, scalable and standardized behavioral tracking could help elucidate impacts of agrochemicals and other stressors in bumblebees, particularly in light of interactive effects that necessitate testing under multiple conditions (Siviter et al., 2021) (Fisher et al., 2023). Second, quantifying how these stress responses vary across species and populations could help bring insight into differential success of *Bombus* species. Bumblebee declines in North America and Europe have been well documented (Cameron et al., 2011), but population trends vary substantially across species (Jackson et al., 2022) and highlight the need for understanding mechanisms underlying species-level differences in behavior and colony performance. In addition, research has focused disproportionately on North American and European *Bombus*. Developing tools that are accessible to researchers in historically underfunded regions could help correct this imbalance and fill critical knowledge gaps in the ecology, evolution, and diversity of social behavior and colony phenotypes of bumblebees.

## Supporting information

Sample BumbleBox Data Output

## Acknowledgments

We thank Rémy Vandame, Alejandra Martínez de Castro, and Andres Castro-Siller for their collaboration in testing and using the earlier iterations of the BumbleBox. We also thank Brett Graham of the Harvard Neurotechnology Core for his assistance in the early stages of development of the BumbleBox software, and to Michael Dillon, Emily Bick, and members of the Crall lab for feedback on initial drafts of this manuscript. This work was supported by USDA National Institute of Food and Agriculture (NIFA) Postdoctoral Fellowship to J.D.C. (grant no. 2019-67012-33596), Agricultural and Food Research Initiative grant no. 2022-67013-36275 from the USDA National Institute of Food and Agriculture to J.D.C., a USDA NIFA Hatch Project (grant no. WIS04062), a Winslow Foundation grant to J.D.C, the Wisconsin Alumni Research Foundation, and Feir Fellowship to AEC.

## Conflict of Interest Declaration

The authors declare no competing interests.

## Supplementary Materials

### Supplementary figures

**Supplemental Figure 1.**
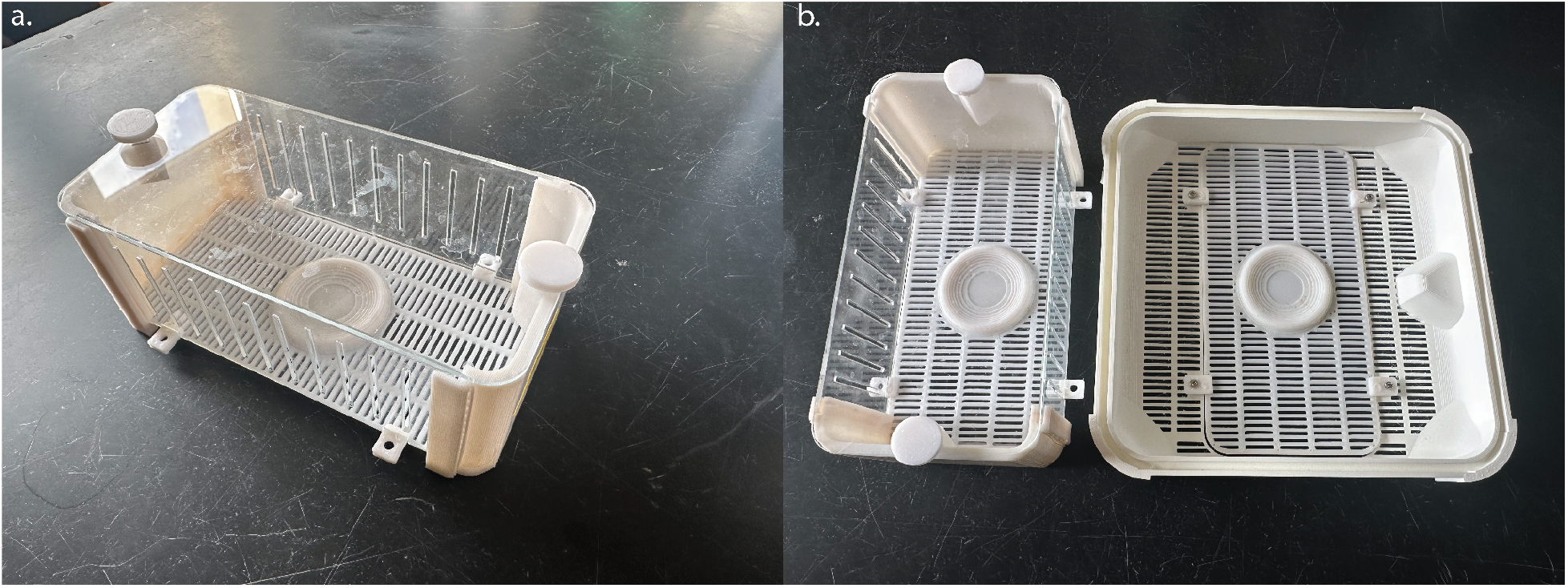
The queen/microcolony starter box. a) The assembled starter box is made entirely from 3D printed and laser-cut acrylic parts. b) A comparison of the starter box and the base of the BumbleBox - the floor of the starter box fits as an insert into the base of the BumbleBox.

**Supplemental Figure 2.**
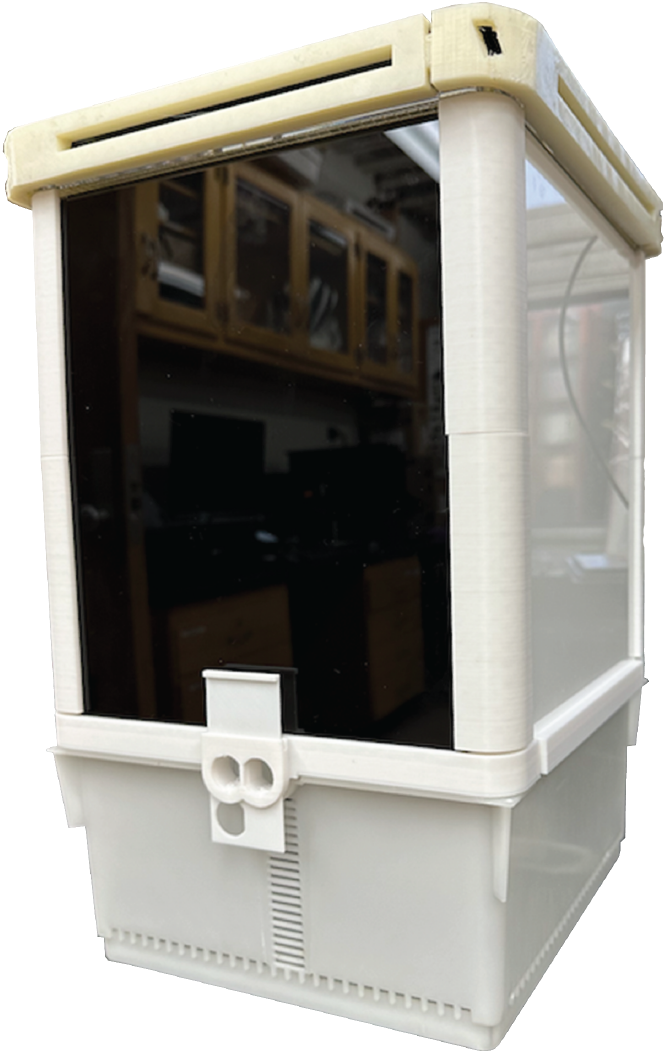
The Koppert-fitted BumbleBox. The BumbleBox design fits on top of a Koppert commercial colony box, extending the height of the box in order to image the entirety of the box floor.

**Supplemental Figure 3.**
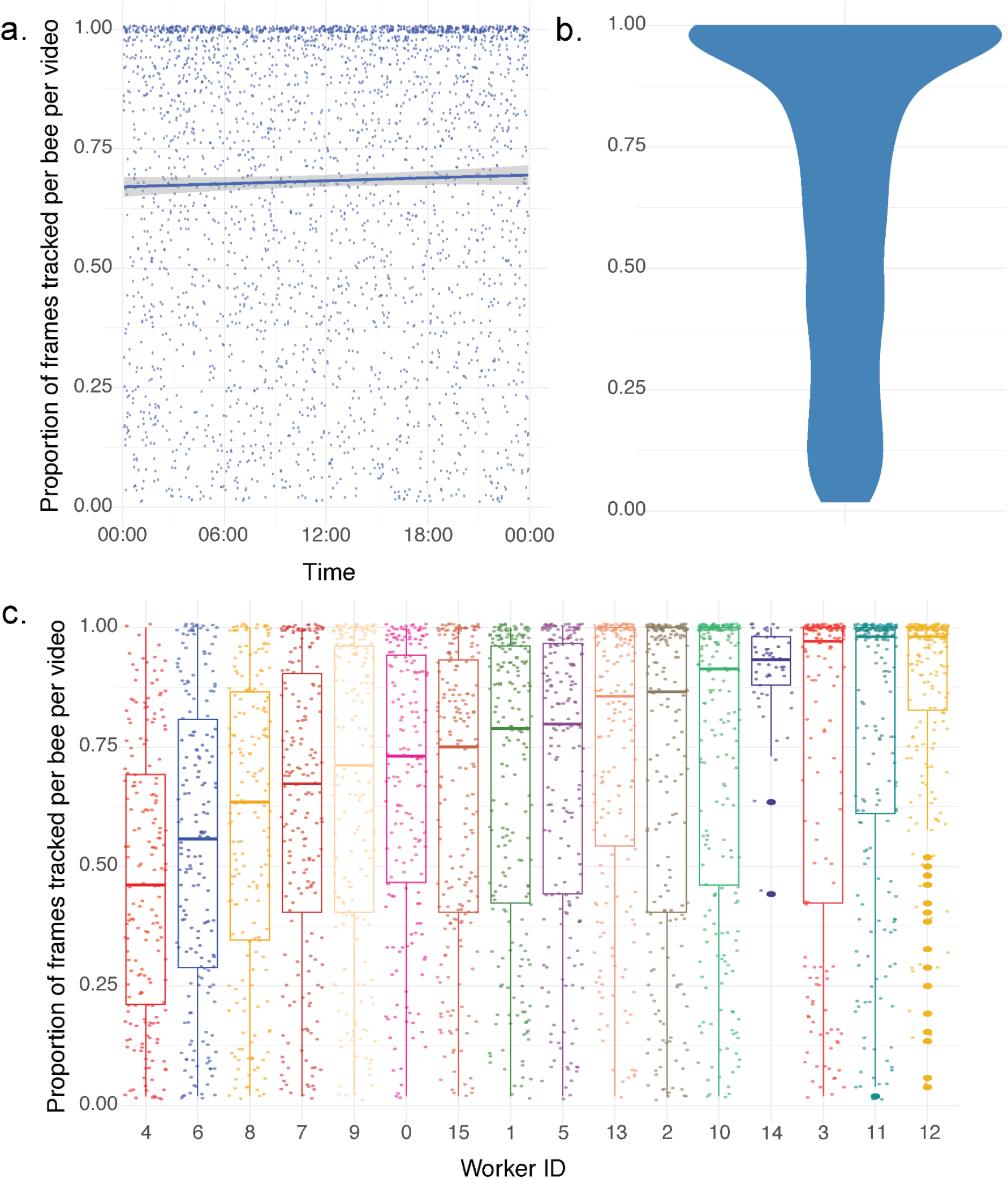
Tracking coverage across 16 workers over 24 hours. a) The proportion of each video each worker is tracked in over time. Each point represents a single worker in a single video, and the LOESS smoothed curve illustrates trends in tracking performance over time. b) Violin plot illustrating the distribution of tracking coverage, across all workers and videos. c) Illustration of the tracking coverage for each individual worker, ordered to increase from left to right by median tracking coverage. Each data point represents one worker in one video, and each boxplot displays the median and interquartile ranges of the data.

## Supplementary text

### Tracking performance analysis, continued from 3.1 of the main manuscript

Out of the 107,235 tracking instances, we found 18 instances of duplicate tags within the same frame, resulting in a duplication rate of 0.017%. Of these, there were only 8 instances where the distance between the x and y coordinates respectively between the two duplicate tags was more than 1 pixel each. In these 10 other instances, the duplication can be considered a double-reading of the same tag. In either case, due to the low rate of duplicate tag reading, the issue is easily resolved by removing both duplicates.

Next, we investigated the proportion of frames that workers were tracked in for each video in which they were tracked in at least one frame, of the 240 videos that make up the dataset.Across all workers, the median tracking coverage was 78.8%, and 25th and 75th percentiles were 42.31% and 98.08%, respectively. Only one individual was tracked in less than 50% of each video it appeared in over half the time (Supplementary Fig 3c).

## Notes

https://github.com/Crall-Lab/BumbleBox

### Competing Interest Statement

The authors have declared no competing interest.

